# Theoretical and practical refinements of sans spike-in quantitative ChIP-seq with application to p300/CBP inhibition

**DOI:** 10.1101/2022.08.09.503331

**Authors:** Bradley M. Dickson, Ariana Kupai, Robert M. Vaughan, Scott B. Rothbart

**Author notes:** Electronic address.

## Abstract

Previously, we introduced an absolute and physical quantitative scale for chromatin immunoprecipitation followed by sequencing. The scale itself was determined directly from measurements routinely made on sequencing samples without additional reagents or spike-ins. We called this approach sans spike-in quantitative ChIP, or siQ-ChIP. In this paper we extend those results in several ways. First, we simplified the calculations defining the quantitative scale. Second, we highlight the normalization constraint implied by the quantitative scale and introduce a new scheme for generating ’tracks’ for siQ-ChIP. We next introduce some whole-genome analyses that are unique to siQ-ChIP which allow us, for example, to project the IP mass onto the genome to evaluate how much of any genomic interval was captured in the IP. We apply these analyses to p300/CBP inhibition and demonstrate that response to inhibition is a function of genomic architecture. In particular, active transcription start sites are only weakly perturbed by p300/CBP inhibition while enhancers are strongly perturbed. Similar observations have been reported in the literature, but without a quantitative scale, those observations have been misinterpreted. We discuss how the siQ-ChIP approach precludes such misinterpretations, which stem from the widespread community practice of treating unquantified and unnormalized ChIP-seq tracks as though they are quantitative.

## INTRODUCTION

The chromatin immunoprecipitation (ChIP) method was introduced in the 1980’s to analyze DNA-protein interactions at specific genomic loci in prokaryotic and eukaryotic cells[1, 2]. In general, the method involves in-cell fixation of chromatin-associated proteins to DNA, chromatin extraction and fragmentation, IP of chromatin fragments with antibodies specific to the target protein or post-translational modification (PTM) state, DNA isolation, and analysis of enriched fragments by hybridization, amplification, and sequencing methods. With few modifications to this method, and recent adaptation for compatibility with high-throughput sequencing (seq), ChIPseq is now widely deployed for studying DNA-associated protein distributions and densities across genomes.[3]

There is a perception that ChIP-seq is not a quantitative method.[4] As such, the chromatin community has developed modifications to ChIP-seq protocols involving the introduction of spike-in reagents of defined concentration at various stages of sample preparation to establish relative quantitative scales.[5–9] The goal of these signal normalization approaches is to enable direct comparison of ChIP-seq results across samples and provide an accurate means of determining, for example, how cellular perturbations impact the distribution of histone PTMs and chromatin-associated proteins across genomes. However, these relative scales are not defined in terms of absolute quantities or units. Moreover, a lack of method standardization and bookkeeping practice makes it impossible to directly compare ChIP-seq datasets from experiment to experiment within the same lab, across different labs, and from datasets compiled as part of large-scale consortium initiatives like the EnCODE Project[10] *even when spike-ins are used*.

We recently introduced sans spike-in quantitative ChIP-seq[11] (siQ-ChIP), a method that emerged from the concept that ChIP-seq is itself inherently quantitative on an absolute scale by virtue of the equilibrium binding reaction in the IP of chromatin fragments. The theoretical model of this equilibrium binding reaction, as introduced in our prior work, proposed that the captured IP mass would follow a sigmoidal isotherm if the reaction was governed by classical mass conservation laws. If we could map the number of sequenced fragments into the total number of fragments contained in the IP product, then we could obtain a quantitative scale through connection to the isotherm. Cellular perturbations that change protein or PTM presentations would emerge as changes in position on the isotherm, and would thus be directly quantitatively comparable.

Informed by continued theoretical analysis and experimental practice, we report here an optimized and simplified crosslinking ChIP protocol for siQ-ChIP. No new reagents or procedures are required for this approach, save for a novel and necessary method to match input chromatin concentrations for IPs. We also present further development of the proportionality constant, *α*, that is needed to compute the siQ-ChIP quantitative scale. The improved expression for *α* is simple to understand, simple to evaluate, and importantly, results in values that are identical between the old and new expression. This new expression reinforces a normalization constraint, ignored by the community, related to how sequenced fragments are aggregated into visual representations. Previously, our analysis was based on three-dimensional representations of sequencing data that obeyed this constraint whereas, below, we introduce a scheme compatible with standard two-dimensional ChIP-seq tracks. This constraint can impact global track shape and has implications on how tracks should be interpreted. We discuss some published misinterpretations as examples. We also introduce two novel modes of automated whole-genome analysis that can be used to easily visualize and compare outcomes of cellular perturbation related to the distribution and abundance of histone PTMs measured by siQ-ChIP.

## MATERIALS AND METHODS

### Cell culture and drug treatment

HeLa cells (ATCC #CCL-2) were maintained in DMEM (Gibco, 11965092) supplemented with 10% FBS (Sigma, F0926) and 1% penicillin-streptomycin (Gibco, 15140122) and were grown in 37^◦^C with 5% CO_2_. Cells were passaged and plated 1 d before drug treatment. Media was then removed and replaced for drug treatment with 10 *µ*M CBP30 (Cayman #14469 Batch: 0473336-73), 10 *µ*M A485 (Cayman #24119 Batch: 0581192-13), or vehicle dimethyl sulfoxide (DMSO) (Sigma 472301, Lot: SHBK2080). DMSO was volume matched for experiments and was either at 0.1% or 0.02% total volume. Cells were treated for 16 h and were then fixed and collected using the protocol described herein.

### ChIP

#### Cell fixation and collection

The volumes listed were for a 10 cm dish of HeLa cells at approximately 70% confluency. Cells were rinsed once with 10 mL of D-PBS (Gibco, 14190136) followed by cross-linking for 5 min in 10 mL of 0.75% formaldehyde (Pierce, 28906) in D-PBS at room temperature. Formaldehyde was removed, and cells were quenched for 5 minutes by addition of 10 mL of 750 mM Tris. Cells were washed twice with 10 mL of D-PBS, scraped into cold D-PBS, collected by centrifugation at 300 g, and snap-frozen in liquid nitrogen. At this point, cells were stored at −80^◦^C.

#### Chromatin Isolation

Cells were then lysed under hypotonic conditions in 1 mL of 20 mM Tris-HCl pH 8, 85 mM KCl, 0.5% NP40 (1 tablet of protease inhibitor (Roche, 11836170001) per 5 mL of buffer) for 30 min on ice. Nuclei (and other insoluble material) were collected by centrifugation at 1300 x g for 5 min at 4^◦^C, lysed by resuspension in 150 *µ*L 50 mM Tris-HCl pH 8, 150 mM NaCl, 2 mM EDTA, 1% NP-40, 0.5% sodium deoxycholate, 0.1% SDS (1 tablet of protease inhibitor per 5 mL of buffer), and passaged five times through a 27-gauge needle (BD #309623 Lot: 0227218). Lysate was then diluted to 500 *µ*L by addition of 350 *µ*L of binding buffer (25 mM HEPES pH 7.5, 100 mM NaCl, 0.1% NP-40). Five *µ*L of RNAse A/T1 (Thermo Scientific, EN0551) was added, and the sample was incubated at 37^◦^C for 25 min. Next, CaCl_2_ was added to a final concentration of 40 mM (21 *µ*L of 1 M) followed by the addition of 75 U (3 *µ*L of 25 U/*µ*L) of micrococcal nuclease (MNase, Worthington Biochemical) and incubated at 37^◦^C for 5 min. MNase was quenched by the addition of 40 mM EDTA (46 *µ*L of 500 mM EDTA), and the total volume was brought to 1 mL by the addition of 425 *µ*L of binding buffer. Next, insolubilities were removed by centrifugation at max speed (about 21,000 x g) at 4^◦^C for 5 min, and the supernatant containing soluble chromatin was collected.

#### Chromatin Measurement

At this stage, 5 *µ*L of chromatin was measured using the Qubit dsDNA HS Assay Kit (Invitrogen, Q32851). To ensure similar chromatin concentrations and match IP conditions, samples were diluted with binding buffer to match each other.

#### Antibody to bead conjugation

For each IP, 25 *µ*L of Protein A coated magnetic beads (Invitrogen, 10008D) were washed once with binding buffer and incubated with either 0, 1.6, 2.5, or 10*µ*L of antibody against the target histone mark. Total volume of bead+antibody was brought to 200 *µ*L using binding buffer and were rotated at room temperature for 15 min. Buffer containing antibody was removed, and beads+antibody were resuspended in 200 *µ*L of soluble chromatin followed by 15 min rotation at room temperature. Fifty *µ*L of chromatin was set aside for input. Unbound chromatin was removed, and beads were vortexed for 10 s with 500 *µ*L of binding buffer. Buffer was removed, and bound material was eluted from beads by vortexing for 10 s in 133 *µ*L of elution buffer (25 mM HEPES pH 7.5, 100 mM NaCl, 1% SDS, and 0.1% NP-40). At this time, the input was brought to 133 *µ*L by the addition of 83 *µ*L elution buffer. Proteins were digested by the addition of proteinase K (Invitrogen, 25530015) to a final concentration of 15 *µ*M overnight at 37^◦^C. The following morning, each DNA sample was purified using MinElute PCR Kit (Qiagen, 28004) and eluted in 30 *µ*L of Buffer EB. Five *µ*L of DNA was quantified by Qubit dsDNA HS Assay Kit. The remaining 25 *µ*L of DNA was frozen at −20^◦^C until it was prepared for sequencing libraries. For comparison of a mark between samples, we performed ChIP of all samples on the same day and made a master mix of bead+antibody for each ChIP target, scaling up all components by the number of samples.

#### DNA gel

DNA fragment size of inputs was checked on 1X TBE 2.5% agarose gels with 1X SYBR Safe (Invitrogen, S33102) to ensure MNase digestion. One *µ*L of NEB ladder (#N3231S Lot: 10047328), run at 60 V for 60 min.

### Library Preparation and Sequencing

Details such as the amount of DNA taken into library preparation can be found in Supplementary Table 1. Library preparation was done using KAPA HyperPrep Kit (Roche, KK8504) with 4 *µ*L of Illumina adapters (IDT, UDI) and sequenced on Illumina NextSeq 500 with a Mid-Output (paired-end 75 bp reads) flow cell. Input libraries had between 83-102M reads, and each IP library had 27-46M reads that passed QC, with 88% of the bases having quality scores ≥30.

### Antibodies list

<u>For ChIP-seq</u>

H3K27ac (Active Motif, 39133 Lot 06921014 – Figure 3), biological replicates from Supplementary Figure SI-Fig. 5A use lots 06921014 and 16119013

H3K18ac (Active Motif, 39755 Lot 26919002) Figure 3 H3K18ac (Invitrogen, MA5-24669 Lot: WB3 187272) Supplementary Figure SI-Fig 5B

<u>For Western Blot SI-Fig. 4</u>

H3K27ac (Active Motif, 39133 Lot 16119013) 1:2000 H3K18ac (Invitrogen, MA5-24669 Lot: WB3 187272) 1:2000

Total H3 (Epicypher, 13-0001 Lot:12320001) 1:50000 Rabbit Secondary (Cytiva, NA934V Lot: 17016966) 1:10000

### NGS Data Processing

We followed exactly the same procedure as previously described[11].

## RESULTS AND DISCUSSION

### A simplified *α*

In this section, we derive a simplified expression for quantitative ChIP-seq scaling. This new expression is more intuitive to understand, easier to evaluate, and more accurate to sequencing outcomes than the previous expression. While values derived from old and new expressions are consistent, the new expression demonstrates a clear and explicit dependence on paired-end sequencing.

In our previous work[11], we built the inherent ChIPseq quantitative scale by first noting that the total number of reads available in a given IP can be written as

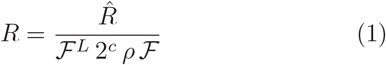

where 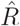 is the total depth of the IP and *R* is the total possible depth if the full IP mass were sequenced. ℱ^*L*^ substituted into *α* to produce is the fraction of library sequenced, *ρ* is the total library concentration divided by the theoretical library concentration (what we call the *library efficiency*), and ℱ is the fraction of IP’d material taken into library prep. The 2^*c*^ accounts for particle doublings encountered during amplification and adaptor ligation. Combining equation (1) with the analogous expression for input, and taking the difference in volumes for IP and input into consideration, one obtains the quantitative ChIP-seq scaling factor *α*

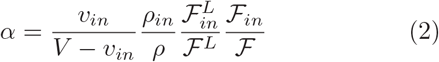

The expression for *α* can be simplified considerably by writing all the factors of *α* in their base units and cancelling as many contributions as possible. Equation (1) can be reduced to (writing all the terms in mass units)

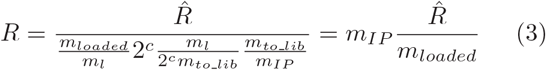

where *m*_*to lib*_ is the mass taken into library prep, *m*_*loaded*_ is the mass loaded onto the sequencer, and *m*_*IP*_ is the full IP mass. The total possible reads that can be extracted from an IP is expressed as the product of the IP mass, *m*_*IP*_, and a reads per unit mass conversion factor 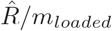. Alternatively, the unitless ratio *m*_*IP*_ */m*_*loaded*_ scales the actual depth 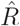 to the total possible depth *R*.

Using equation (3), and the analogous result for input reads, we can rewrite *α* in a more intuitive and simplied way, where

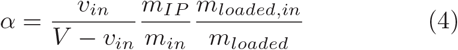

Finally, the fraction 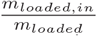 can be reinterpreted through the following observation. The total sequencing reads generated by a single flow-cell are commonly split among several samples. Loading a multiplexed flowcell can be idealized as: Each sample is standardized to the same molarity, then different volumes are taken from each sample and pooled. The volume fraction of each sample now corresponds to the fraction of total particles that come from that sample. In this circumstance, the fraction of the flow-cell’s reads that will be consumed by each sample is given by its volume fraction in the pool. The expectation is that the conversion from moles of chromatin fragments to sequencer reads is constant, 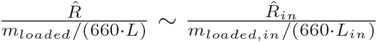. Here, 660 is the average molecular weight of a DNA base pair (*g/mol/bp*), *L*_*in*_ and *L* are the average fragment lengths for input and IP respectively. These are library fragment lengths, reported in our case by a Bioanalyzer. The moles to reads relationship between input and IP can be rearranged and substituted into α to produce

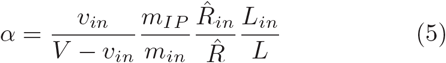

The symbols 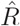 and 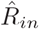 represent the number of sequencing reads (or fragments) generated by IP and input, respectively. We have cancelled the factors of 660 *g/mol/bp* in equation (5). Figure 1 shows the correspondence between equations (2) and (5) for the data reported in this paper. The new *α* can be evaluated for expected depth or actual depth, where as the previous form explicitly used mass loaded into sequencing and is therefore limited to expected depth. Figure 1 clearly demonstrates the correspondence of equations (2) and (5) when expected depth is used. The new *α* is simpler to understand, easier to evaluate, and more accurate to sequencing outcomes because we can make use of the obtained depth rather than the requested depth (which was previously encoded through ℱ^*L*^ and 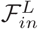).

**FIG. 1:**
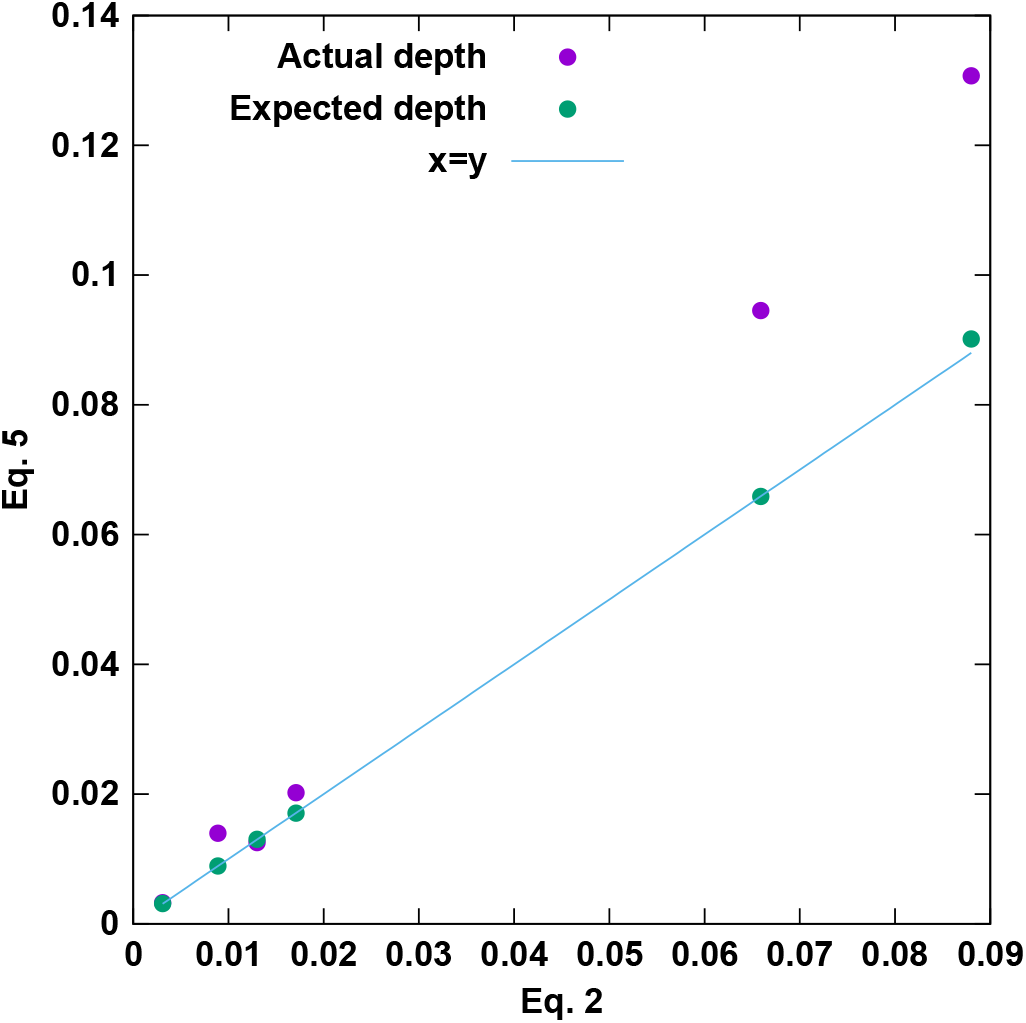
Direct comparison of Equations (2) and (5) where depths are taken as actual (observed) or expected.

The most informative perspective on *α* comes by viewing it as the ratio of two factors 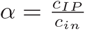 with

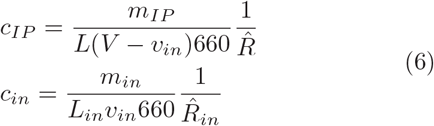

Each of these coefficients is written as the product of two quotients. The first expresses the IP or input mass as a concentration by direct units conversion. The second is a normalization factor. Therefore, if *f* (*x*) is a browser track of the IP sequenced fragments, which is just a histogram of fragments intersecting base pair *x*, then *c*_*IP*_ *f* (*x*) is the concentration of DNA that overlaps *x* that was bound in the IP reaction. This projection of bulk concentration to genomic location is valid if, and only if, 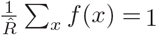. When the track *f* (*x*) is built, each fragment can be counted only so that 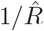 normalizes *f* (*x*).

This normalization constraint was respected by the three-dimensional efficiency we previously introduced (see Eq. 2 of Reference 11 where *ê*(*x, L*) plays the role of *f* (*x*)). However, genome browsers are not designed to present three dimensional data, *e*.*g*., fragment position, length, and capture efficiency. A further complication is that the standard process of building tracks for use in a browser yields tracks that do not satisfy this normalization constraint. If, for example, the *i*-th sequenced fragment accumulates a +1 at every base pair that it intersects, then the *i*-th fragment is over-counted *L*_*i*_ times, with *L*_*i*_ the length of the fragment in base pairs. Note that paired-end sequencing is required to correctly determine the *L*_*i*_.

Accumulating +1*/L*_*i*_ at each intersected base pair, instead of +1, resolves all of the above concerns entirely. A track built this way is a proper histogram and is normalized by the number of observations that went into the histogram, 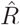 for an IP and 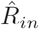 for input, and is suitable for genome browsers. In this scheme, each base pair in a fragment is equally weighted, just like when +1 is accumulated. However, different fragments are not equally weighted unless they have the same length. In particular, longer fragments will effectively contribute with lower weight because there is a greater uncertainty in ’where’ the important binding event was when that fragment was captured.

In Figure 2, we illustrate six sequenced fragments and show the outcome of building a track using the +1 or +1*/L*_*i*_ accumulations. The set of six fragments form two ’islands,’ where fragments within an island do not overlap fragments from outside the ’island.’ When +1*/L*_*i*_ is used, each ’island’ of piled fragments will reflect a sum of fractions that looks like 1*/L*_1_ + 1*/L*_2_ + …. Each ’island,’ then, can be understood as having its own common denominator for the summation of the fractions. In our example, the left island has a common denominator of 30 while the right island has a common denominator of 56. The fact that islands have different common denominators allows the islands to have different final scales. For example, when the +1 weights are used, the left and right islands form peaks of equal height (Figure 2B). When the +1*/L*_*i*_ weights are used, the right island forms a shorter peak than the left island (Figure 2C). This is because the right island is comprised of longer fragments and these fragments convey a larger uncertainty about where the peak ought to focus.

**FIG. 2:**
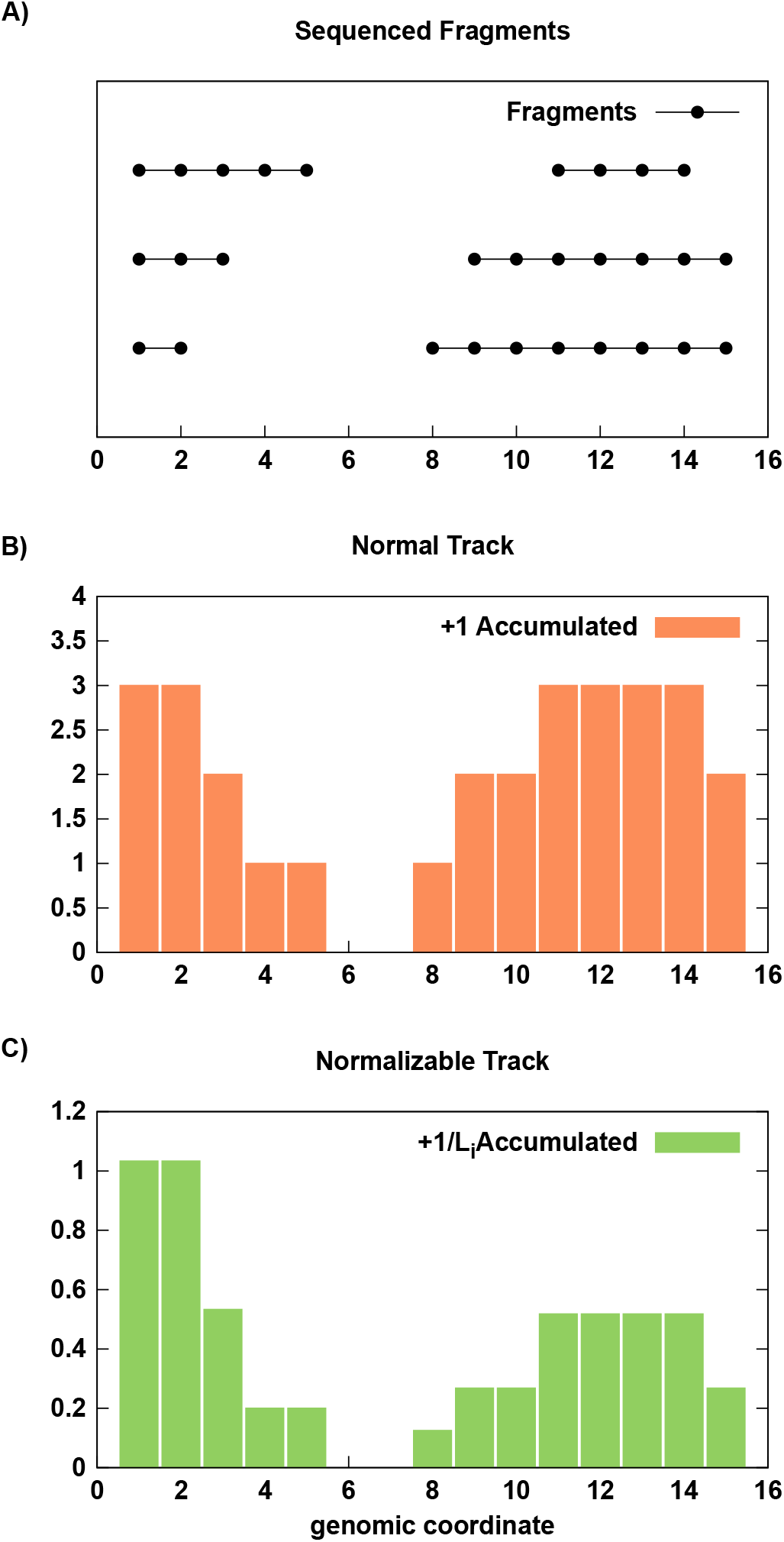
Example track builds using 6 sequenced fragments (A) and accumulating +1 (B) or +1*/L*_*i*_ (C) when counting fragments at each genomic position.

The scaling of the sequencing tracks by *α* (or *c*_*IP*_, etc) can only be correctly interpreted as a projection of the physical IP outcome onto genomic position if 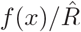 is a normalized probability distribution. This constraint on building *f* (*x*) is a key insight that has not been provided by any other analysis of ChIP-seq, yet it is critical for preservation of physical scale, which as we have just demonstrated, can impact track shape across the genome. In light of this constraint, arbitrary scaling rules like RPKM (reads per kilobase of million mapped) become unnecessary. Moreover, any material quantity can be projected onto the genome with a correctly assembled track. For example the IP mass itself can be projected onto genomic coordinates, 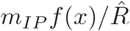, allowing one to compute the mass contributed to the IP from any genomic interval.

To summarize, the quantity *c*_*IP*_ *f* (*x*) is an estimate of the concentration of chromatin bound in the IP that originated from position *x*. Likewise, *c*_*in*_*f*_*in*_(*x*) is an estimate of the total concentration of chromatin in the IP reaction that originated from position *x*. The siQ-ChIP “track” in quantitative units is given by 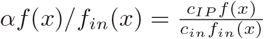 and is an estimate of the IP binding efficiency at position *x, i*.*e*., the fraction of chromatin originating from *x* that is bound in IP.

The above development of *α* and its dependence on average fragment length motivate some comments on our practice of chromatin fragmentation. Complex distributions of fragment length introduce error in mass-to-concentration conversions and may artificially inflate IP capture masses. We found Micrococcal Nuclease (MNase) digestion of chromatin produced narrow fragment length distributions, especially when compared to the typical outcome of sonication. Notably, and consistent with prior work[12], MNase digestion does not lead to bias in the ability to observe heterochromatic nucleosomes with this assay protocol.

Finally, we note that this simplification of *α* requires the practitioner to report 6 parameters at the start of compiling siQ-ChIP data. The previous form of *α* required dozens of entries to compute the same value. It is also worth noting that all of the following analysis is automated in the current version of the siQ-ChIP software, which can be found on GitHub.[13]

### CBP/p300 inhibition via CBP30 and A485

To demonstrate the utility of siQ-ChIP with working examples, we considered the impact of inhibiting p300/CBP on acetylation at lysines 18 and 27 on histone 3 (H3K18ac and H3K27ac, respectively). We consider the effects of two inhibitors, CBP30[14] and A485[15]. CBP30 targets the bromodomain of p300/CBP while A485 targets the acetyltransferase domain. This system has been recently characterized by others[16–19], allowing a comparison of our analysis of ChIP-seq to existing results. Importantly, all of the previous work reports and interprets ChIP-seq data that is not quantified in absolute terms. We therefore directly examine the role of absolute quantification in interpreting ChIP-seq observed consequences of the two modes of p300/CBP inhibition.

Central to the siQ-ChIP paradigm is the antibody:chromatin isotherm. This isotherm is, in reality, a many dimensional surface, with the coordinates in its domain being the concentration of antibody, and the concentrations of all epitopes. For each of our experimental contexts (CBP30 inhibition, A485 inhibition, DMSO control) we determined the antibody:chromatin isotherm by titrating antibody. The isotherms are shown in Figure 3, where all epitope coordinates are held fixed and the antibody concentration was titrated. Because the total chromatin concentration is fixed (to within experimental ability), the change in isotherm as a function of target-epitope concentration is approximated by the changes seen in moving from DMSO to CBP30 to A485. Keep in mind that the total chromatin concentration is fixed, only the concentration of p300/CBP dependent epitopes has changed. Figure 3 informs on the full surface of the antibody:chromatin isotherm and is a useful global landmark for assessing consistency of repeat experiments. IP masses should always be reported with ChIP-seq data, along with evidence that the IP conditions were matched (*e*.*g*., by reporting input masses). The IP mass can confer notions of losses or gains which can be used as a reference against the apparent changes observed in sequenced data. The power of these isotherms is that we have more than one observation on the mass at more than one antibody load. These observations can be considered for self-consistency as well as referenced against any future repeats.

**FIG. 3:**
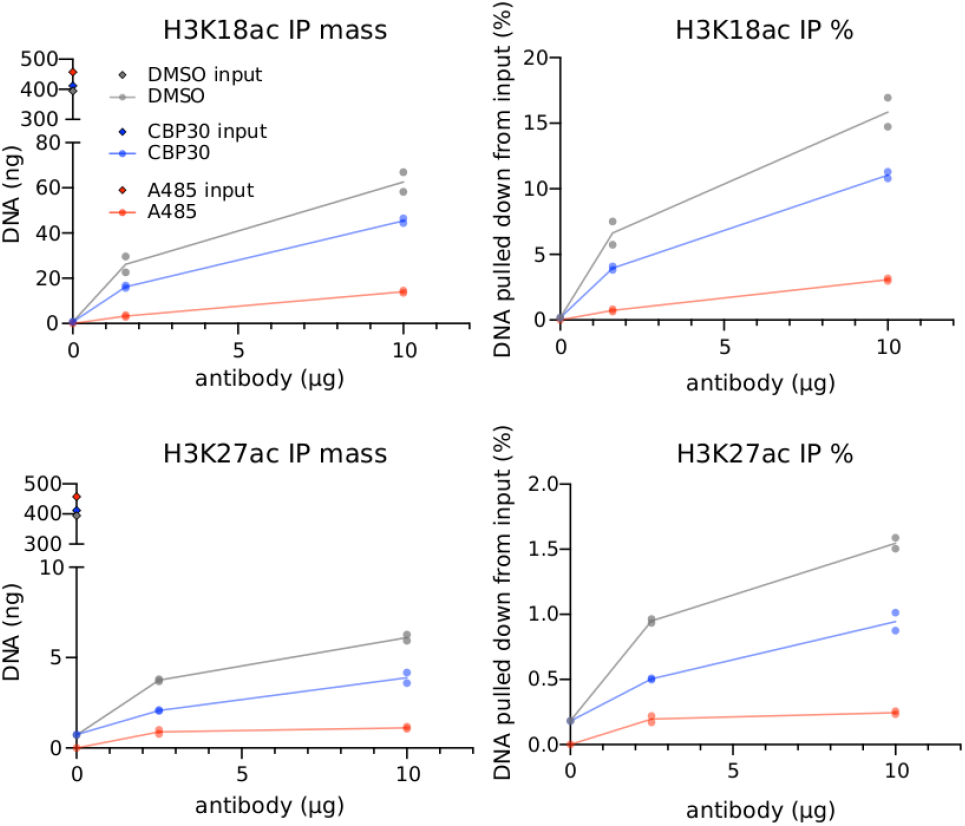
H3K18ac (top) and H3K27ac (bottom) ChIP antibody titrations for HeLa chromatin extracted following DMSO, CBP30, or A485 treatment. Data from two independent experiments are reported as total mass and percent input.

Several observations come from the data presented in Figure 3. First, H3K18ac produces substantially higher IP masses than does H3K27ac. Both the H3K18ac and H3K27ac antibodies appeared to have their isotherm inflection points to the left of the 2.5 and 1.6 *µ*g antibody loads, evidenced by the fact that both antibodies appear to be approaching saturation — Each antibody displayed diminishing returns after an ∼ 8 fold increase in concentration. The loads 1.6 and 2.5 *µ*g differ between antibodies because of limiting reagent constraints. Figure 3 suggests either that H3K18ac is more abundant in HeLa chromatin than H3K27ac or that the H3K18ac antibody is binding to many off-target chromatin species. This data does not argue that the H3K27ac antibody is weaker, and thereby captures less mass, because of the change in slope from 0 to 2.5 and from 2.5 to 10 *µ*g antibody. A weaker antibody will not simply plateau at a lower captured mass but will instead plateau only at higher antibody loads. Reports coming from mass spectrometry suggest that H3K18ac is more abundant than H3K27ac[20, 21], consistent with the isotherms in Figure 3.

Second, the isotherms show that A485 is effective (more-so than CBP30) at globally reducing levels of H3K18ac and H3K27ac with the treatment paradigm used here. At low antibody load, CBP30 produced a 1.4-fold reduction in IP mass for H3K18ac and a 1.9-fold reduction in IP mass for H3K27ac. The moderate effects of CBP30 on these PTMs is consistent with prior work.[17] Meanwhile, under the same treatment and IP conditions, A485 produced a 6.1-fold reduction in H3K18ac IP mass and a 4.9-fold reduction in H3K27ac IP mass. We carried the 1.6 and 2.5 *µ*g antibody points into sequencing, because this is where we would predict the highest antibody specificity [11].

These isotherms give us the most critical parameters in equation (5), the input and IP masses. The remaining contributions to *α* are the input and IP reaction volumes, and the average fragment length and depth which are determined during sequencing. The following sections address analysis of the sequencing data.

### Whole genome analysis using annotations

The formal model of the IP binding process at the heart of siQ-ChIP explicitly expands chromatin into a list of all possible chromatin modification states[11], where we call each state a species. Below we explore the possibility that genomic annotations may provide a means of grouping several distinct species into larger classes. Each annotation is, of course, the aggregate of several species, but decomposition of the IP products into annotations turns out to be very useful nonetheless. This analysis provides a simple way to understand how the global distribution of fragments is changed by experimental perturbation.

We take the 15 state model put forward by the NIH Roadmap Epigenomics Consortium as our model for chromatin-state (and therefore epitope/species) labels [22]. To estimate the distribution of the sequenced fragments with respect to the 15 distinct annotations, let *a*_*i*_, with *i* = 1, 2, …, 15, indicate the *i*th annotated state. Let *f*_*IP*_ (*a*_*i*_) be the total number of IP reads that intersect any genomic interval annotated by *a*_*i*_ (if a fragment intersects twLo or more, arbitrarily take the first one). Notice that 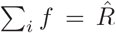, just as we required above. As a novel representation of the sequencing data, the IP mass can be projected onto the annotations as *L*(*V* − *v*_*in*_)660 × *c*_*IP*_ *f*_*IP*_. Figure 4A shows the IP masses from H3K27ac and H3K18ac projected onto the annotations, thus decomposing the total IP masses into contributions from each class of genomic annotation. Each column of Figure 4A sums to the total mass for that experiment.

**FIG. 4:**
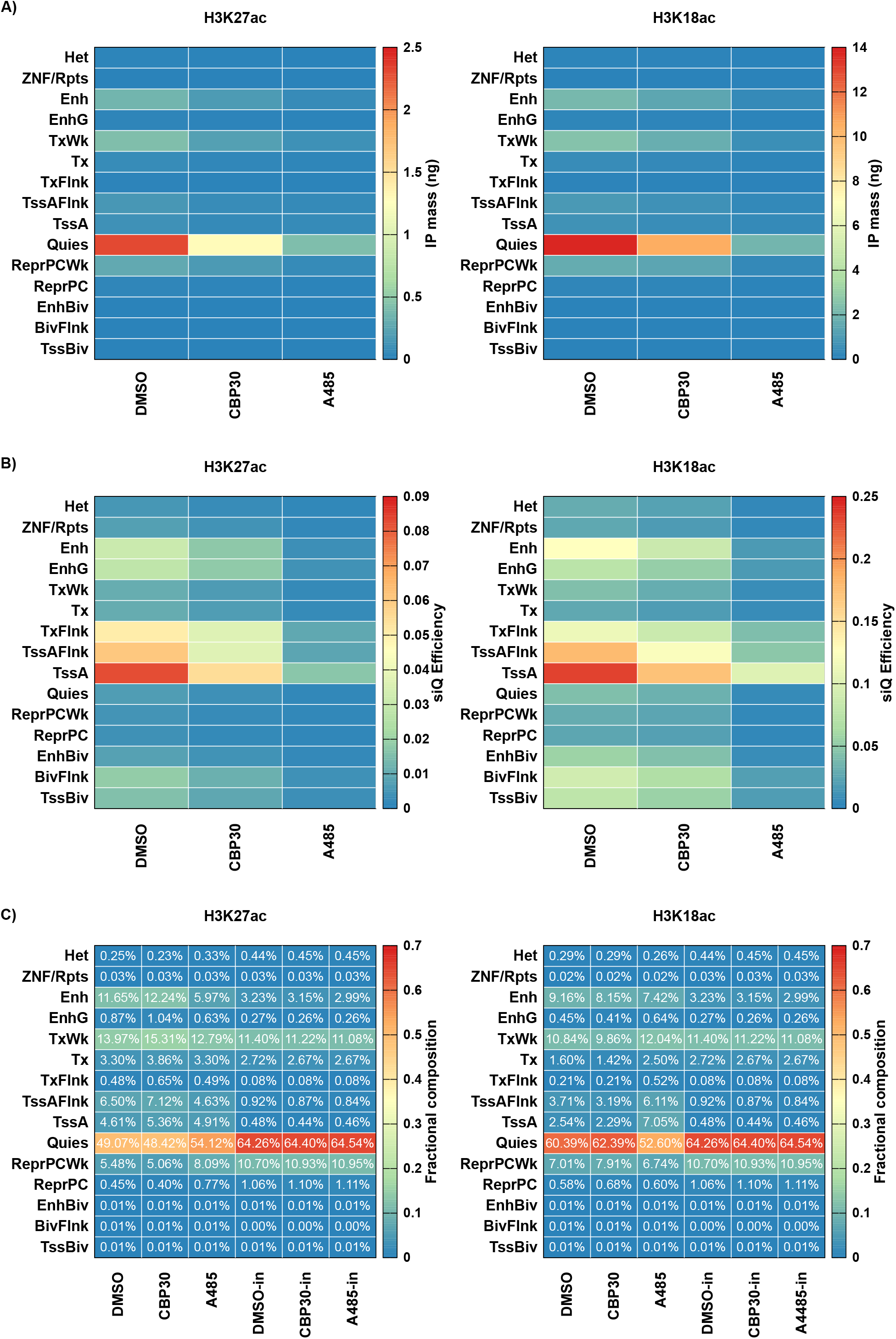
(A) Mass of IPs projected onto annotation after DMSO, CBP30, or A485 treatment. (B) siQ-ChIP capture efficiency. (C) The fractional composition of IPs.

Figure 4B reports the siQ capture efficiency (*αf*_*IP*_ */f*_*in*_) of H3K18ac and H3K27ac for each annotation. Notice that the largest IP mass in either H3K18ac or H3K27ac was due to the Quies annotation (Figure 5A), but the largest capture efficiency was due to the TssA annotation for both H3K18 and H3K27 (Figure 4B). The capture efficiency of Quies is actually small for both H3K18ac and H3K27ac. The IP masses in Figure 4A do not report on the input composition, but the siQ capture efficiency does. This is why quantification in terms of capture efficiency is important and more meaningful than mass alone. Likewise, the mass itself does not report on *enrichment*. By looking at capture efficiency, we can see that both antibodies enrich for annotations that reflect active chromatin states. On the other hand, looking at the masses shows that a great deal of sequencing reads were consumed by Quies regions and that, surprisingly, the capture at Quies is dependent on p300/CBP activity.

**FIG. 5:**
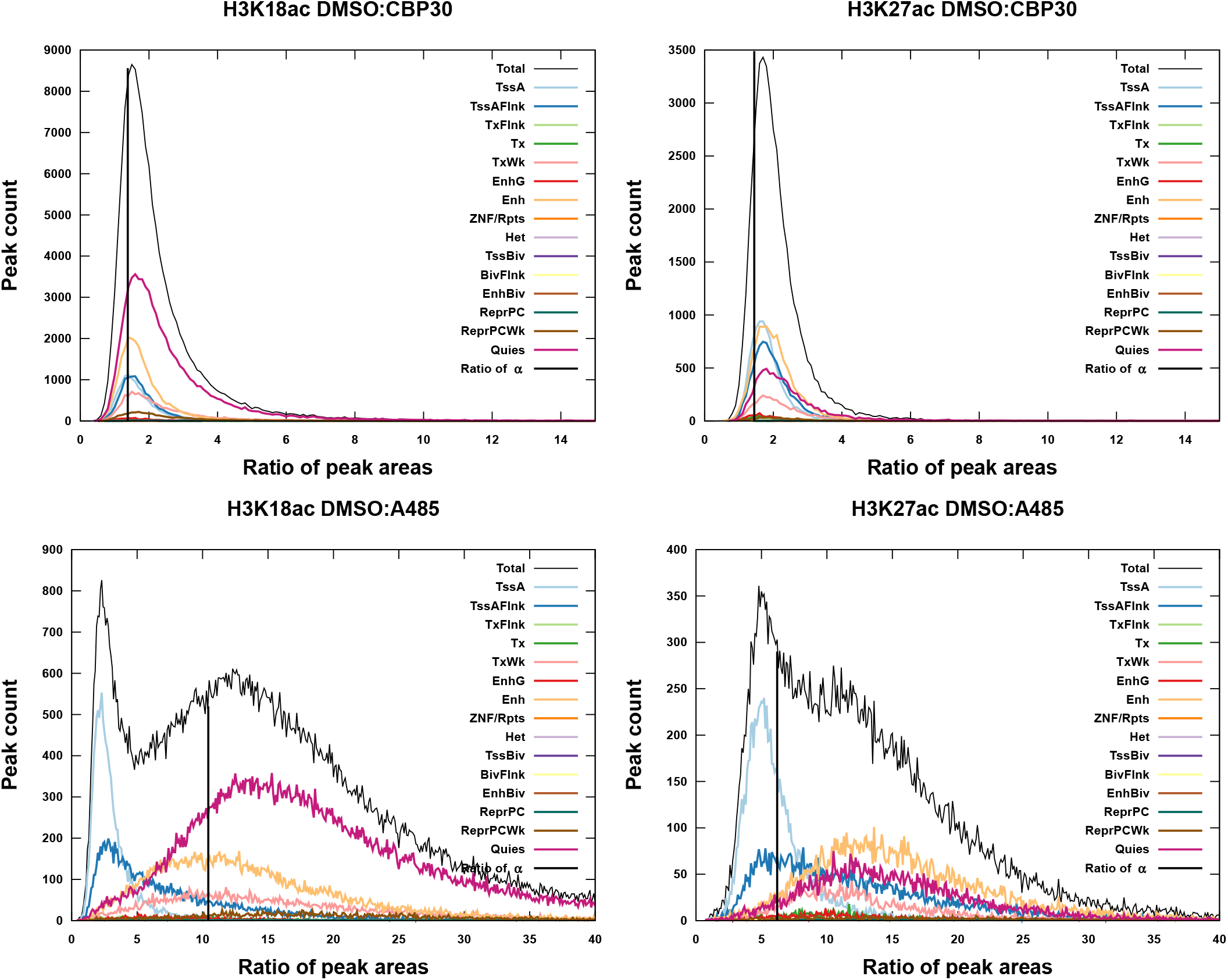
The distributions of peak responses to p300/CBP inhibition, *µ*(*r*), in sequencing peaks. The response distribution is shown as a total and decomposed into contributions by annotation.

Along with siQ-ChIP, we previously introduced the *fractional composition* of the IP as a way to present the distribution of IP products, and we studied this distribution through simulations to show that it can behave in some counterintuitive ways.[11] The fractional composition is identical to the distribution of fragments over the annotations, 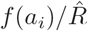. The use of annotations as a proxy for species allows us to examine the fractional composition of actual IPs, and to visualize how the distribution of IP products responded to different p300/CBP inhibition paradigms. Figure 4C shows the fractional composition for each IP and input, computed as 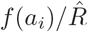. Perhaps the most notable feature emerging from this analysis is the increase in TssA- and TssAFlnk- (active promoter flanking) associated fragments in the H3K18ac pulldown after A485 treatment. (2.8- and 1.6-fold, respectively, Figure 4C) This increase reflects the increased probability of observing a TssA associated fragment, when randomly selecting fragments from the IP. This increase does not indicate an increase of H3K18ac PTM at TssA annotations, as both the capture efficiency and captured mass are down after inhibition (Figure 4A,B).

The fractional composition shows us that the IP product distribution is reshaped by A485, which in turn suggests that the H3K18ac antibody is still preferentially binding chromatin fragments with TssA annotations. There is either residual H3K18ac in these annotations or there is a significant off-target species recognized by the H3K18ac antibody. We did not observe drastic reshaping for the H3K27ac antibody.

To summarize the whole genome analysis based on the histogram of annotations (*f* (*a*_*i*_)), which does not involve making genome browser tracks nor calling peaks, p300/CBP inhibition via A485 results in a deep loss of IP mass and capture efficiency for both H3K27ac and H3K18ac antibodies across all annotations. The H3K18ac antibody in particular shows a residual enrichment of TssA annotations after A485 treatment (Figure 4C), while the overall capture of those annotations is impaired (Figure 4A and B). The TssA, TssAFlnk and TxFlnk annotations incurred the weakest losses, while Enh (enhancer) annotations were severely impacted according to both antibodies. We note that GCN5/KAT2A is associated with TssA genomic intervals[23] and has been shown to have H3K18ac and H3K27ac activity *in vitro* [24]. A hypothesis consistent with all of these observations is that GCN5/KAT2A, which is not inhibited by A485, is maintaining some level of H3K18ac/K27ac at TssA, TssAFlnk, and TxFlnk but not at Enh.

Interestingly, the largest single mass component of the IPs are Quies annotations. This annotation responded significantly to A485 inhibition through both antibodies, suggesting p300/CBP is active in these regions of the genome. Moreover, the lack of focused p300/CBP activity in Quies is consistent with the hypothesis that these regions act as a sink[25] for excess p300/CBP activity, allowing the accumulation of a non-functioning reservoir of acetylation for recycling[26]. Inhibition of p300/CBP through the inhibitor CBP30 showed modest mass and efficiency losses but demonstrated no significant reshaping of the IP-product distribution for either antibody. We did not try to improve CBP30 impacts by altering treatment paradigm. It remains to be seen whether CBP30 can drive a response similar to that of A485 with an optimized treatment.

### Whole genome analysis using browser tracks

In this section, we describe how siQ-ChIP tracks are computed, aggregated into a database of peak-wise comparisons, and how this database can be used to quickly obtain whole genome conclusions about the data. The database records all genomic intervals corresponding to track peaks as well as several quantitative attributes. The database itself is not a record of a single track, but rather a record of comparisons between tracks. As discussed above, all tracks are made with the +1*/L*_*i*_ accumulation rule. Details of generating siQ quantified tracks and detecting peaks are given in Supplementary Information (SI-Fig. 1, 2, and 3 and associated text).

In general, a most common use of ChIP-seq is to test how the ChIP-seq signal reacts to experimental perturbation. To do this here, we first identify an interval X in a control track and then investigate that same interval in an experimental track. For our p300/CBP inhibition experiments, this means the DMSO track was taken as a control and either CBP30 or A485 data was taken as the experimental track. For each interval detected in the control track, the area under the signal *s*(*x*) is computed for both control and experimental tracks. Supplementary Information gives details on building *s*(*x*) = *αf* (*x*)*/f*_*in*_(*x*) and identifying the complete set of *𝒳*_*i*_ generated by IP. The minimal Fréchet distance[27] between the two tracks on the interval is also computed, which provides a numerical assessment of how similar the two tracks are in shape within the given interval. This shape information is included in the database of peaks, but we do not make much use of it here because the degree to which shape is reproducible is currently unstudied. Application of metrics like the Fréchet distance will allow future study of shape. SI-FIG. 2 illustrates this metric with current data, and all drug treatments are summarized in SI-Fig. 3.

The final database is a list of the intervals, the area under *s*(*x*) on the interval for experimental and control tracks, the difference in shape between the tracks, and a few other attributes as noted in the documentation of our tools.[13] Once this database is built, several modes of analyses parallel to those shown in Figure 4 are possible.

A most informative analysis is represented in Figure 4B, where the siQ-efficiency of capture is shown as a function of genomic annotation. Of central importance in ChIP-seq is how the sequencing signal, namely the peaks, changed due to experimental perturbation. We define the Response on interval *𝒳*_*i*_ between *s*_*cntr*_ and *s*_*exp*_, a control and experimental track respectively, as

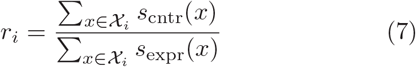

The sums in numerator and denominator represent the area under *s*(*x*) on the interval *𝒳*. The response quantifies the change in area under a peak upon experimental perturbation.

A whole-genome characterization of this response is possible by looking at the distribution of responses *µ*(*r*). This distribution (unnormalized) is shown for both A485 and CBP30 inhibition in Figure 5. The x-axis in these Figures, *r*, is the ratio of areas as DMSO:A485 or DMSO:CBP30. The y-axis is the number of peaks that had a response of *r* ± *dr* with *dr* a binwidth. The results after CBP30 inhibition are again not striking. This has been evident since the isotherm was determined (Figure 3). On the other hand, the results characterizing A485 inhibition show not only a strong response but also a bimodality of response.

The total response distribution can be deconvoluted into contributions from the different annotations as

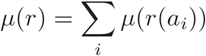

In practice, this amounts to grouping peaks by the annotation they fall on. This deconvolution is shown in Figure 5. Of particular interest is the distribution of responses for the TssA annotations. For both H3K27ac and H3K18ac, the distribution *µ*(*r*(TssA)) is shifted to the left and has a long tail to the right side. For H3K18ac, the maximum in *µ*(*r*(TssA)) is near *r* = 2 and shows that most of these peaks have small changes in area after A485 treatment compared to DMSO. These are peaks that respond weakly to A485. The long right-side tail indicates that there are still many peaks that did respond to A485 and had a loss in area. For H3K27ac, we found the maximum in *µ*(*r*(TssA)) at *r* = 5 meaning there is typically a five-fold reduction in area after A485 treatment. This response is smaller than expected (10-fold for H3K18ac and 6-fold for H3K27ac), where the expected response is estimated as the ratio of *α*’s for the two experiments. We conclude that the response is less than expected for TssA annotations in both H3K18ac and H3K27ac, with the response being severly muted in H3K18ac data. Additionally, there is a larger response in shape perturbations for Enh than TssA (SI-Fig. 3).

Figure 5 combined with Figure 4 indicates which genomic features/annotations respond to perturbation, how significant that response is, and whether there are peaks associated with the response. Thus, without looking at a single browser track, we have completely described the results of p300/CBP inhibition across the entire genome. The shape of signal in the whole genome can be understood through this analysis without ever focusing our attention on a single isolated peak or gene. To connect this abstract and general characterization to the more familiar representation of tracks, in Figure 6 we show a region of the genome where one can appreciate both responsive and non-responsive peaks. One may also appreciate that H3K27ac has a more muted response on TssA in this window, and H3K18ac has nearly no response on TssA in this window. Meanwhile, peaks on Enh annotations are lost. The diversity of peak responses summarized in Figure 5 are clearly visible even in this small window on chr1.

**FIG. 6:**
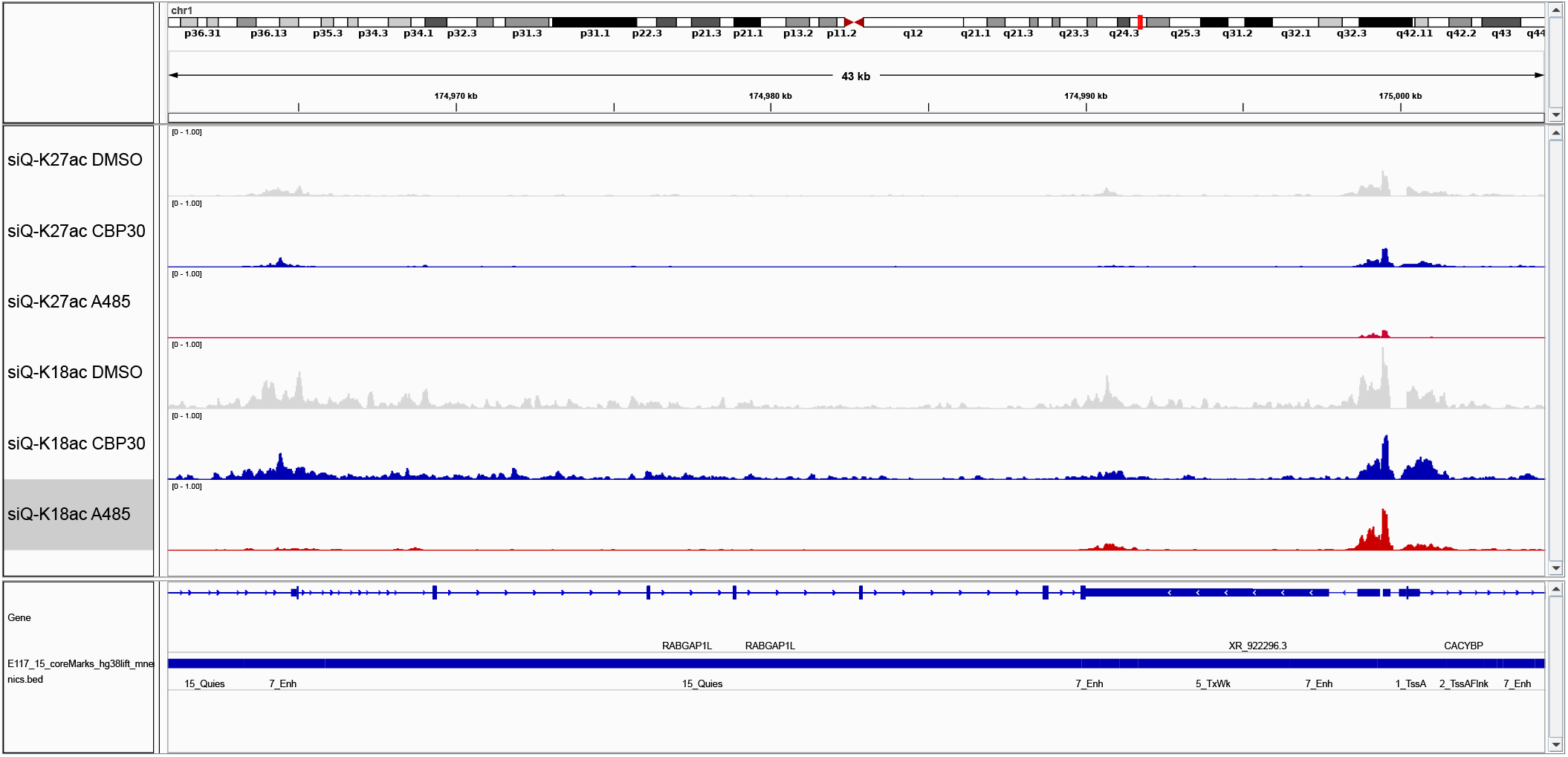
A sample browser shot showing diversity of TssA and Enh response in H3K27ac and H3K18ac data. All ChIP-seq data ranges are [0, 1] for direct comparison between tracks and conditions. The window coordinates are chr1:174960844-175004779.

### Quantitative scale and ChIP-seq interpretation

Figure 4C shows that there is an increased fraction of TssA associated fragments in the H3K18ac IP after A485 treatment, and Figure 5 shows that the peaks associated with TssA annotations have an extremely weak (roughly 2-fold) response. siQ-ChIP applies a global scaling factor, *α*, to the sequencing tracks, and the H3K18ac *α* was decreased 10-fold upon A485 treatment. For peaks in the sequencing track to decrease by only 2-fold while *α* decreases by 10-fold, peaks in the unscale track (before *α* is applied) must have increased in magnitude by 4 to 5-fold. This increase offsets the decrease in *α*, leading to the muted response. This very clearly illustrates the problem of treating unscaled ChIP-seq data as though it were quantitative, where the physical scale of the IP binding reaction is ignored.

Figure 7 shows metaplots for H3K27ac and H3K18ac at TssA and Enh. In light of everything discussed above, the metaplots in unscaled units (indicated as IP/input) should be interpreted as reflecting the ratio of probability of finding fragments at a position (IP to input), not the *amount* of PTM. H3K27ac shows no evidence of redistribution in the (unscaled) metaplots for TssA, consistent with the analysis above. H3K27ac displays a loss in probability on Enh, again consistent with Figure 4C. We clearly see a large increase in track height at TssA for H3K18ac in unscaled units (Figure 7B), with little change at Enh. In all cases the scaled version of these plots reflect the significant losses that were indicated in Figure 4B.

**FIG. 7:**
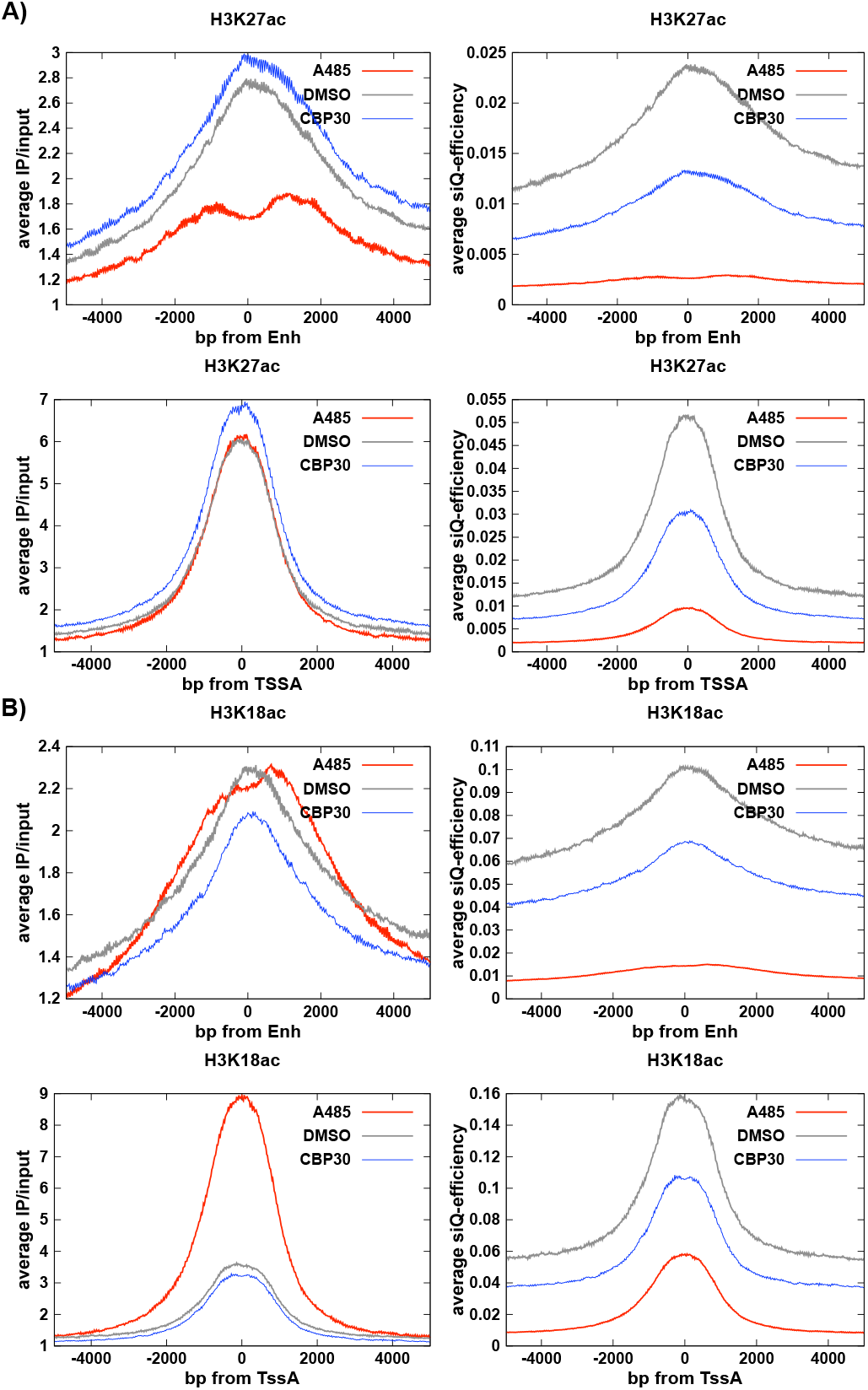
Metaplots in unscaled (left) and siQ-scaled (right) units for Enh and TssA annotations captured by (A) H3K27ac or (B) H3K18ac antibodies.

The problem with unscaled ChIP-seq is that an easy misinterpretation is to conclude that there is an increase in H3K18ac at TssA after A485 treatment. Indeed, this conclusion is reported in several papers.[16–19] In one study, the increase in H3K18ac signal at TssA was described as an ’increase in the average level of H3K18ac and H3K27ac.’[28] In another study, using spike-ins, it was ’observed that the H3K27ac mark was reduced at [TssA] whereas H3K18ac was slightly increased.’[17] Yet another recent report observed that ’surprisingly, only 226 [of 807] downregulated genes exhibited hypoacetylation of H3K27 under the condition of CBP/p300 HAT inhibition.’[19] Consistently, treating unscaled ChIP-seq data as though it were quantitative leads to the conclusion that the targeted acetylation may be unaffected or even increased in some genomic regions — counter to expectations and contrary to the indication of global loss in the isotherms (IP mass capture, Figure 3), which mirror what is seen by other global measures like western blots (Figure 3a of Reference 28 and SI-Fig. 4).

The emergent constraint that our sequencing tracks must sum to the total depth means that when we normalize to the depth, the resulting track is interpreted as a probability. An unscaled track that respects this constraint expresses the probability that a random fragment contains any given base pair. Thus, for the H3K18ac antibody, it is only *more probable* that IP’d fragments are associated with TssA after A485 treatment. The sequenced tracks are *f* (*x*) in the above discussion, and thus in general are representative of the probabilities of finding coverage of genomic intervals — even if the normalization constraint is not enforced. An increased probability is not an indication of an increase in absolute amount of PTM. For the most part, this misinterpretation does not do much harm, aside from the errant notion that the PTM has actually increased, especially when orthogonal data are integrated. However, using the correct physical scale will not only avoid seemingly inconsistent conclusions, it may also help bring ChIP-seq and RNA-seq, along with other orthogonal observations, into better alignment.[19] We speculate that the homeostatic mechanism[28] implied by the increased probability of finding H3K18ac fragments at TssA after A485 exposure is GCN5/KAT2A. Indeed, we speculate that this alternate and independent route to H3K18ac at TssA likely drives the missinterpretation in all the above-cited p300/CBP perturbation studies.

## CONCLUSIONS

We have described an intuitive simplification of the siQ-ChIP scaling factor *α*, introduced a sensitive ChIPseq protocol for use with crosslinked samples, and developed a genome-wide pipeline for siQ-ChIP data analysis that allows one to easily visualize the distribution of IP mass across annotation classes, and to investigate the full distribution of responses elicited by any cellular perturbations. We hope these advances improve the acceptance and applicability of the siQ-ChIP quantitative method.

Additionally, our protocol gives a simple and controlled distribution of fragment lengths because we use MNase rather than sonication. This improves the accuracy of any mass-to-concentration units conversions where the average fragment length is used.

We have also shown there is a strict condition on how the sequenced data can be used, a condition forbidding over-counting of fragments. This may seem like a minor point but it leads to the interpretation of siQ-ChIP data as a mass distribution, which shows how mass, or concentration of captured species, are distributed along the target genome. Given numerous schemes that have been developed to normalize sequenced data, our proposal to avoid over-counting and normalize solely to the depth to yield 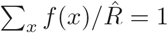 is novel. Interpretation of sequencing results as generating a proper probability distribution precludes the common misinterpretation of sequencing results as quantitative. The constraint on overcounting and the interpretation of quantified ChIP-seq as a ’mass distribution’ make a compelling argument for the soundness of the siQ-ChIP method and suggests an incompleteness of other quantification schemes. This perspective also reveals that spike-in normalization is simply the ratio of two probability functions (the sequencing track of interest and the ”track” or count of fragments of spikein) where at least one of the functions is built with an improper overcounting of reads/fragments.

It is worth discussing that even after *α* and the siQ-ChIP protocol are streamlined, some practitioners will not find this approach to be easier than standard ChIPseq. Moving from qualitative to quantitative experimentation does require additional diligence and care, and there is a learning curve. Because siQ-ChIP is quantitative, there are several practical details that should be mastered and validated. These details include checking that antibody is captured on the magnetic beads used in the IP, minimizing complexity in the chromatin fragmetation products, minimizing bead-only capture in the absence of antibody without invoking any additional practices of bead blocking or pre-clearing. Not to mention, siQ-ChIP dictates that IP conditions be matched and we provide one method of doing just that. Interestingly, even if one uses spike-ins for ChIP-seq the IP conditions should be matched. Yet, no evidence or mention of matching is ever included in publication. The protocol above covers all of these concerns and represents a significant step toward standardizing a quantitative ChIP-seq practice.

All codes for analysis and figures are available at GitHub[13], as are the full database files for all peaks in Figure 5. The genomics data are available via GEO code GSE207783.

## AUTHOR CONTRIBUTIONS

B. M. D., A. K., and R. M. V. data acquisition; B. M. D. software; B. M. D. analysis; B. M. D. and S. B. R. supervision; B. M. D., A. K., and R. M. V. validation; B. M. D. and S. B. R. investigation; B. M. D. visualization; B. M. D., A. K., R. M. V. methodology; B. M. D. and S. B. R. writing-original draft;B. M. D. project administration; B. M. D. and S. B. R. writing-review and editing; S. B. R. and R. M. V. funding acquisition.

## Supplementary Information

### ALTERNATE DERIVATION OF *α*

The siQ-ChIP scale *α* can be obtained as a units conversion applied to the IP reaction efficiency as follows. The heart of siQ-ChIP is the realization that the IP is subject to the basic mass conservation laws that govern all reversible binding reactions. Namely, the total antibody concentration is equal to the sum of the free antibody and bound antibody concentrations. Because of this, the IP mass must follow a sigmoidal form, where increasing antibody concentration causes increased IP mass up until the reaction is saturated. As we explain next, the work of siQ-ChIP is concerned with two features: the determination of the isotherm and the units conversion that maps IP mass to concentration of antibody-chromatin complex. The concentration of complex is what sets the quantitative scale for siQ-ChIP.

In more formal terms, the sum of free antibody (*AB*^*f*^) and bound antibody takes the following form

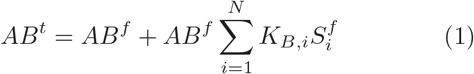

where we used the traditional binding constant definition 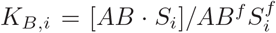. *S*_*i*_ is the *i*-th species or epitope that interacts with the antibody and [*AB* · *S*_*i*_] is the concentration of complex. The total antibody mass is also subject to a conservation of mass constraint for each species, 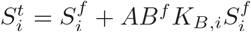 where 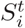 is the total concentration of species *i*. 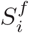 is the free (or unbound) concentration of species *i*.

The symbol 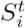 represents the concentration of a chromatin ’state’. Without trying to enumerate all possibilities, these could include all mono-nucleosome fragments that present a defined set of histone modifications. There may be another species 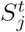 for the di-nucleosome fragments that present the same modifications. Yet another term, 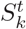, for mono-nucleosomes presenting different modifications or combinations of modifications, and so on.

Of interest here is the solution to these mass conservation laws. The solution is just the set of values 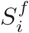 and *AB*^*f*^ that would simultaneously satisfy all of the conservation equations. If we knew the binding constants *K*_*B,i*_ then we could generate the solution numerically. Of course, we do not know the binding constants and we also don’t know how to enumerate all of the terms in the conservation laws, but we have a very handy way to make these shortcomings moot: We determine the actual IP mass empirically, which is itself the sum of all the bound fragments whatever they are and however they came to be there. We can empirically determine this correct mass without needing to know all the terms and constants exactly.

Formally, we have the total bound concentration of chromatin 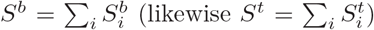, which for our model can be expressed as 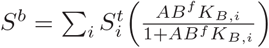 where we used the bound concentration 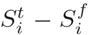. The total *S*^*b*^ is the sum of sigmoids thus, as described above, we anticipate that *S*^*b*^ will plateau or saturate when *AB*^*t*^ is increased.

The key for siQ-ChIP is that this concentration *S*^*b*^ can be converted to mass using the average molecular weight per base pair (660 g/mol/bp) and the average fragment length *L*, yielding *m*_*IP*_ = *S*^*b*^(*V* − *v*_*in*_)660*L* which is the IP mass. The factor (*V* − *v*_*in*_)660*L* converts from concentration units to mass units. Now, once *m*_*IP*_ is determined empirically *S*^*b*^ can be estimated using this unit conversion. We can do a similar thing with the determined input mass, *m*_*input*_ = *v*_*in*_660*L*_*in*_*S*^*t*^ where *S*^*t*^ is the total chromatin concentration. The quantitative scale put forward by siQ-ChIP is based on the fact that the total IP capture efficiency can be expressed as 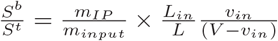.

Because some of the IP and input masses will be sequenced, we have knowledge of the genomic coordinates for a representative collection of the chromatin fragments. Using *x* to denote genomic coordinates and *f* ^′^(*x*) to denote any proper summary of the sequenced fragments (*e*.*g*. 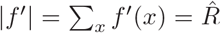 with 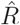 the sequencing depth), we can state that 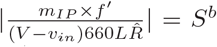. We say *proper* here because 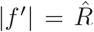 implies that no fragment can be counted more than once. This places a strict constraint on how sequencing tracks are built and interpreted. Typical practice will over count sequenced fragments, with each fragment counted once for each base pair in the fragment. The key result here is that for *m*_*IP*_ *f* ^′^(*x*) the conversion factor 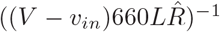 projects *m*_*IP*_ *f* ^′^(*x*) to *S*^*b*^(*x*), which is an estimate of the concentration of bound fragments at *x*.

**SI-Fig. 1:**
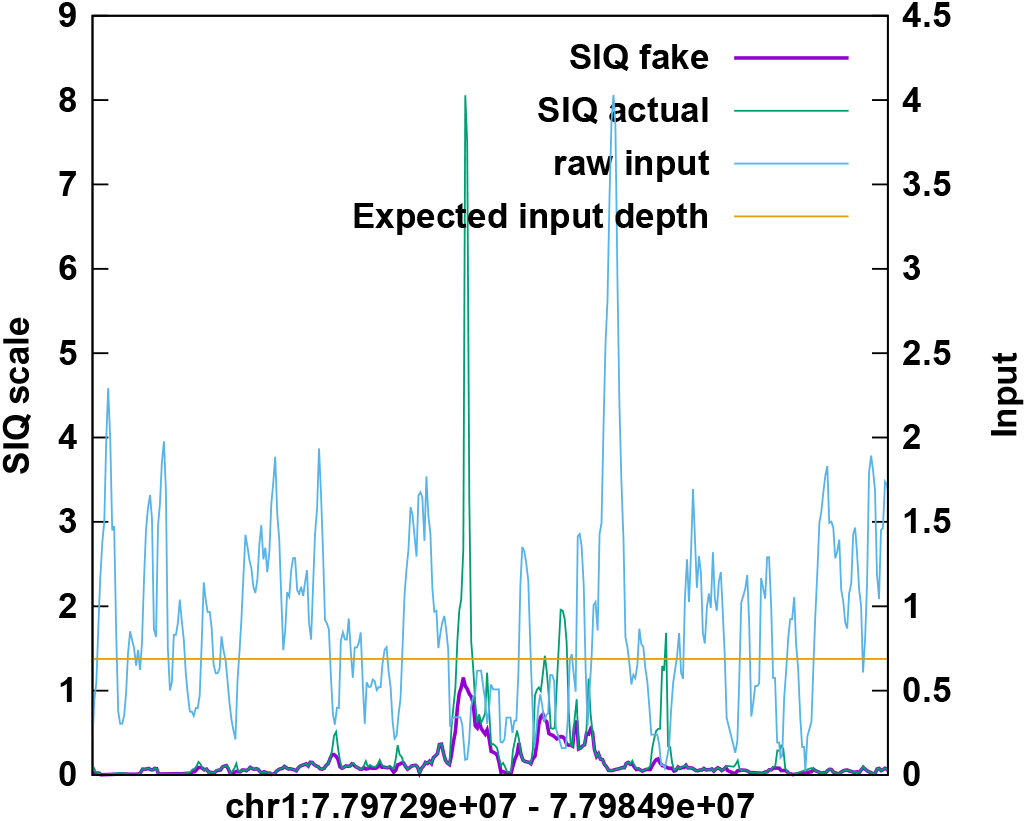
The impacts of using expected input, or ’fake input’, to regularize siQ-scaling.

### PROCESSING SEQUENCING DATA

The siQ-ChIP scale is built on the IP to input ratio because it expresses efficiency of capture. Ultimately, this leaves us to evaluate *αf*_*IP*_ (*x*)*/f*_*in*_(*x*) and to deal with the inevitable case that *f*_*in*_(*x*) ∼ 0 while *f*_*IP*_ (*x*) *>* 0. In these cases the IP demonstrates that the genomic region represented by *x* was present in the chromatin but for statistical reasons has not been presented in the input sequence data.

Because the sequenced input fragments are expected to be binomially distributed along the genome, we estimate the average expected depth, ⟨*d*⟩, of input at any position *x* as 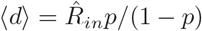 where *p* is the probablity of hitting any base pair in the genome. In our case we use bins larger than a single base pair and *p* is adjusted to this width. (*p* = 30*/*3200000000 for bins of 30 base pair and a total of 3200000000 bases.) Any time *f*_*in*_(*x*) *<* ⟨*d ⟩* we replace the input with ⟨*d* ⟩. We refer to this replacement as ’fake input’ and an example of how this impacts data is shown in SI-Fig 1. The siQ-scale should not be larger than unity for any reason other than noise in the determination of *α*. SI-Fig 1 shows how using the ’fake input’ resolves the over unity problem, where it results from sampling errors in the input track. Over unity peaks are still possible, but are less likely.

The siQ-ChIP sequencing track is given by *s*(*x*) = *αf*_*IP*_ (*x*)*/f*_*in*_(*x*). To call peaks in *s*(*x*) we first compute

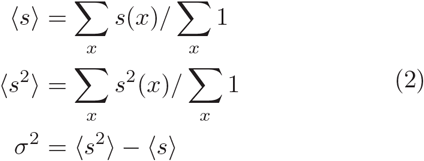

Any genomic interval *𝒳* that has signal satisfying *s*(*x*) > ⟨*s*⟩ for all *x* ∈ *𝒳* and *s*(*x*) > ⟨*s*⟩ + 3*σ* for some *x* ∈ *𝒳* is understood as displaying a peak. This is a simple choice for selecting intervals that have signal larger than apparent background, we did not experiment with values other than 3*σ* but one can set this value in the siQ-ChIP scripts.

As noted in the Main text, part of the database of peaks includes the Fréchet distance between control and experimental data. The Fréchet distance is a metric of shape-similarity between a peak in the control track and the experimental track. This similarity is computed for each interval X, where the tracks on the interval are mapped to the unit square. We map to the unit square so that there is no unit based disparity between the x- and y-coordinates of the tracks and so that the notion of shape is independent of the height of the peaks. To appreciate the quantitative shape comparison, one can imagine the unit square as a visual display, like a projector screen, comprised of pixels. If the control and experimental tracks are ploted on the display, (*dF*)^−2^ gives us an idea of the most pixels the display can have while still allowing the two curves to look similar by eye. A large number of pixels implies a high resolution match, corresponding to a small *dF* value. Conversely, low resolution matches have large values of *dF*.

For example, a value of *dF* = 0.2 gives us 25 pixels while a distance of 0.4 gives us a 6 pixel display. This small displacement of 0.2 in the value of *dF* generates a 4-fold reduction the effective resolution for comparing the data. As a rough guide, values smaller than 0.3 will be generally agreed upon as looking similar where larger values will not. In SI-Fig 2 we illustrate how the metric looks for several actual peak comparisons.

The extent to which peak shapes ought to be conserved between samples or treatments has not been quantitatively characterized. SI-Fig 3 reports all the shape response distributions. We point out that the units and scale of the Fréchet metric take some getting used to. To help calibrate to the scale of *dF*, SI-Fig. 2 reports on a few values of *dF*.

SI-Fig. 4 reports our global observations of histone acetylation after p300/CBP inhibition. Global losses are clearly reported for A485 while little can be appreciated for effects of CBP30.

SI-Fig. 5 reports on a biological repeat of the isotherms for chromatin:antibody reactoins and reports all bead-only capture amounts. No bead-blocking or preclearing is used, and almost all bead-only capture masses are below 1% by mass.

**SI-Fig. 2:**
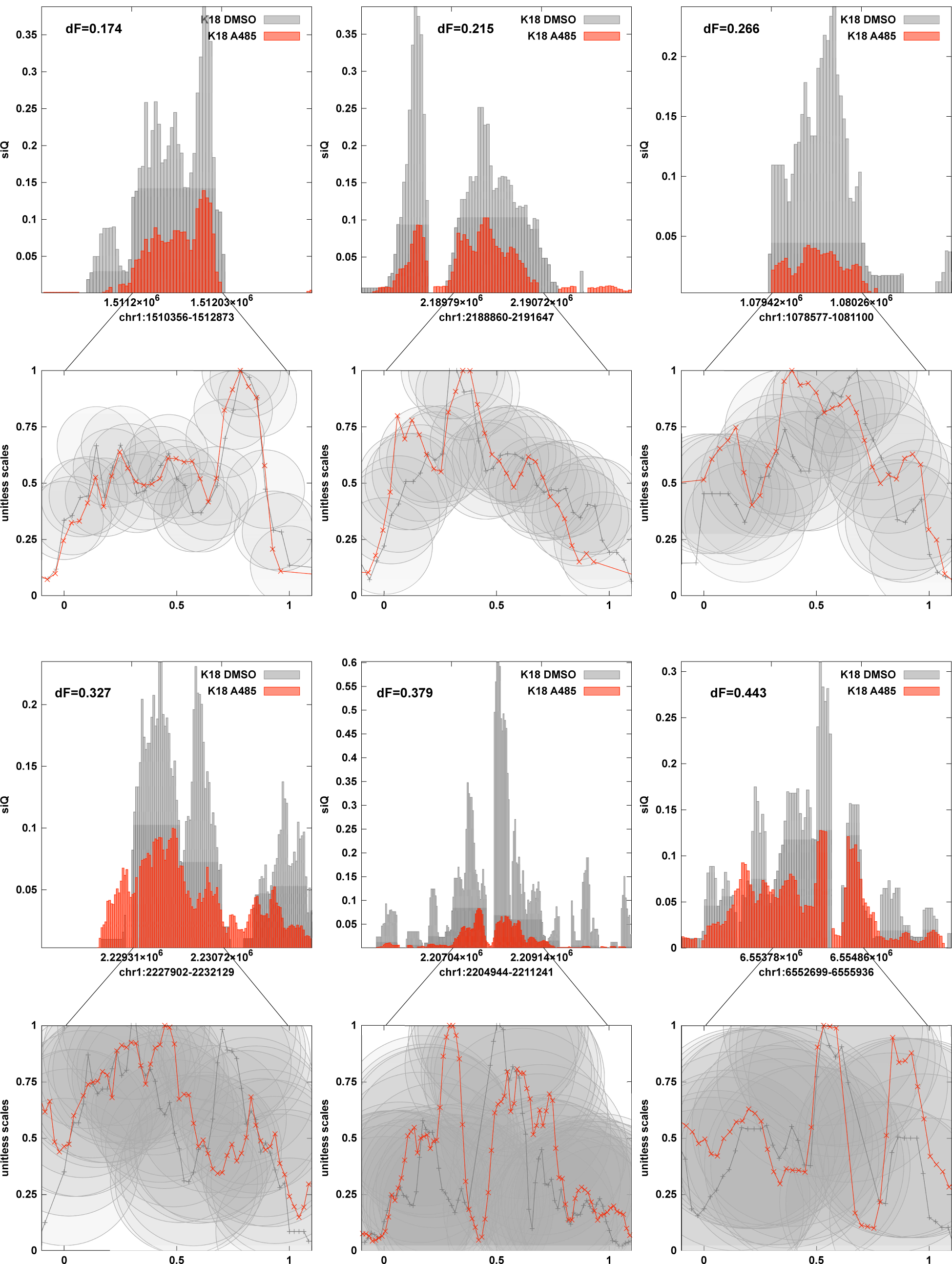
Examples of peaks in a comparison of DMSO and A485 tracks with the K18ac antibody. The peaks are projected to a unitless rectangle and the Fréchet distance is computed. The circles have radii matching the Fréchet distance *dF*. The interval containing a peak in the control track is marked by tics on the top of each bar graph and is expanded to the unitless rectangle where the Fréchet distance is illustrated. The region called to contain a peak runs from x=0 to x=1, in each case, as indicated.

**SI-Fig. 3:**
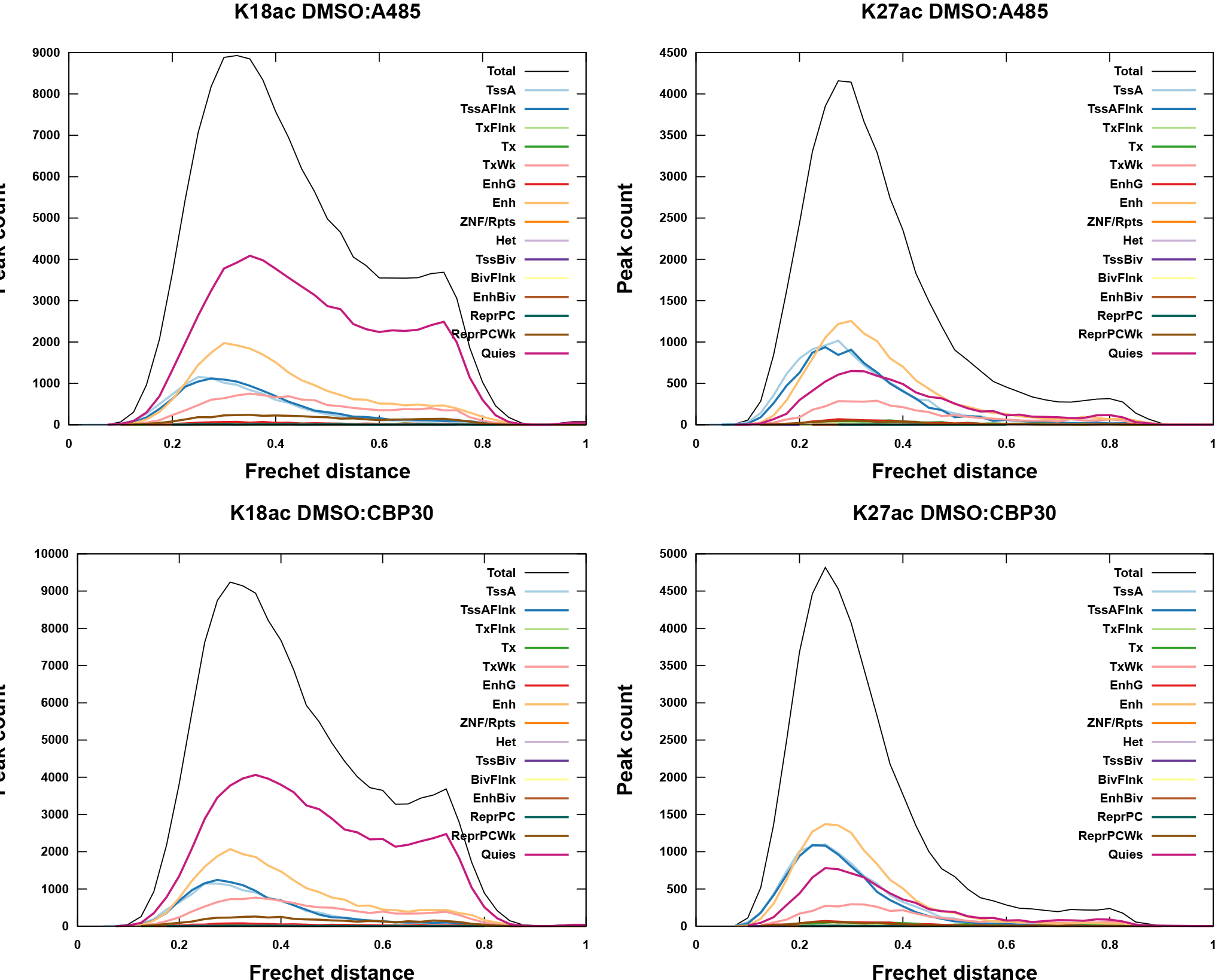
Fréchet response distributions for all drug treatments.

**SI-Fig. 4:**
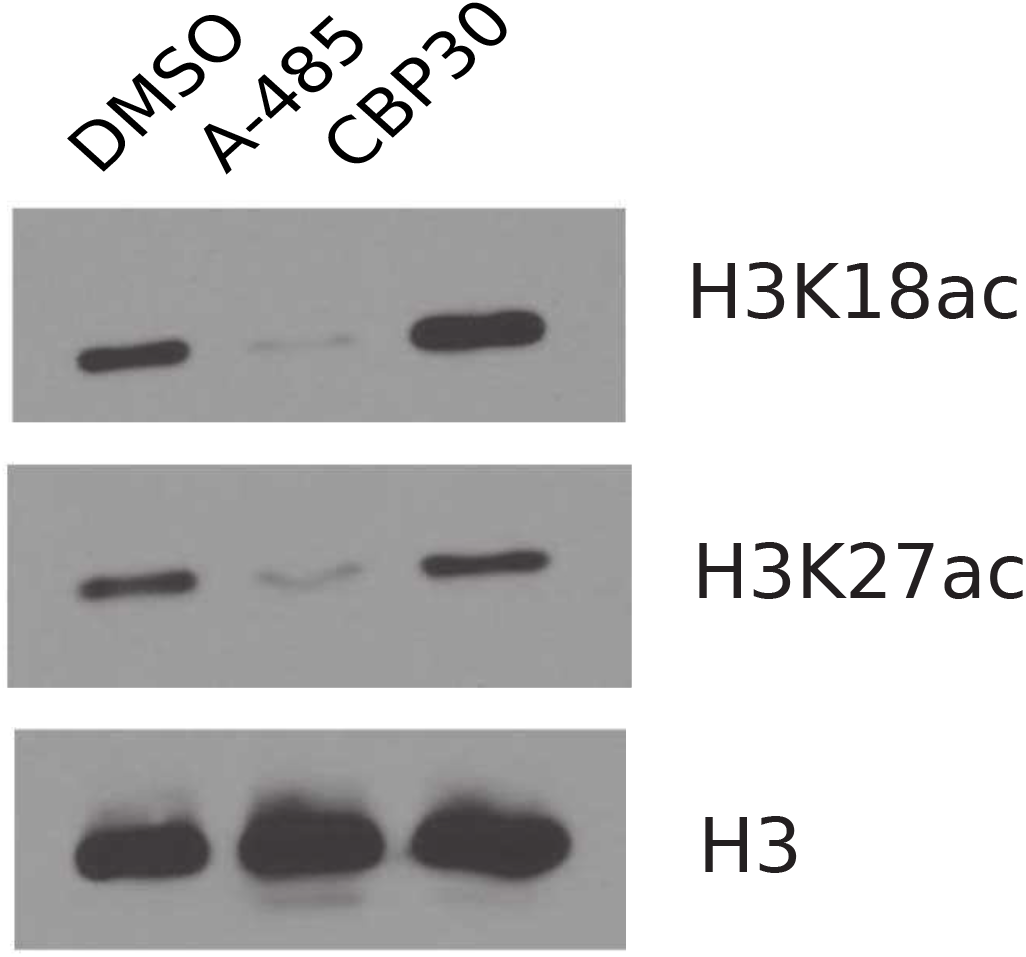
Western blot for H3K18ac or H3K27ac in treated HeLa cells.

**SI-Fig. 5:**
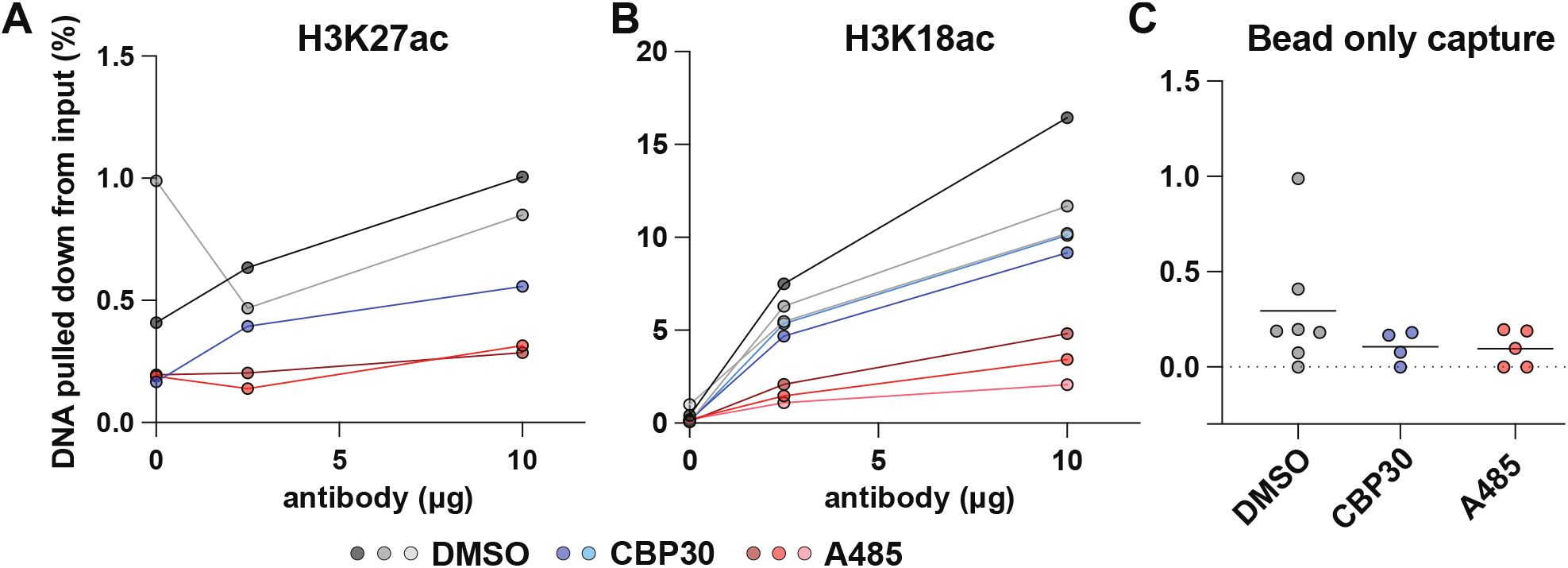
Biological repeats of (A) H3K27ac isotherms, (B) H3K18ac isotherms, and (C) bead-only capture as percent mass. Note that this H3K18ac repeat is with antibody: Invitrogen, MA5-24669 Lot: WB3 187272

